# A complex and dynamic redox network regulating oxygen reduction at photosystem I

**DOI:** 10.1101/2023.09.28.559979

**Authors:** Umama Hani, Belen Naranjo, Ginga Shimakawa, Christophe Espinasse, Hélène Vanacker, Pierre Sétif, Eevi Rintamäki, Emmanuelle Issakidis-Bourguet, Anja Krieger-Liszkay

## Abstract

Thiol-dependent redox regulations of enzyme activities play a central role in regulating photosynthesis. Beside the regulation of metabolic pathways, alternative electron transport has been shown to be subjected to thiol-dependent regulation. We investigated the regulation of O_2_ reduction at photosystem I. The level of O_2_ reduction in leaves and isolated thylakoid membranes depends on the photoperiod in which plants are grown. We used a set of Arabidopsis mutant plants affected in the stromal, membrane and lumenal thiol network to study the redox protein partners involved in regulating O_2_ reduction. Light-dependent O_2_ reduction was determined in leaves and in thylakoids of plants grown in short day and long day conditions using a spin-trapping EPR assay. In wild type samples from short day, ROS generation was twice the amount of that in samples from long day, while this difference was abolished in several redoxin mutants. An in vitro reconstitution assays showed that thioredoxin m, NADPH-dependent reductase C (NTRC) and NADPH are required for high O_2_ reduction levels in long day thylakoids. Using isolated photosystem I, we also show that reduction of a PSI protein is responsible for the increase in O_2_ reduction. Furthermore, differences in the membrane localization of thioredoxins m and 2-Cys peroxiredoxin were demonstrated between thylakoids of short day and long day plants. Finally, we propose a model of redox regulation of O_2_ reduction according to the reduction power of the stroma and the capabilities of the different thiol-containing proteins to form a network of redox interactions.

## Introduction

Thiol-dependent redox regulations of enzyme activities play a central role in regulating photosynthesis. It has been well established that key enzymes of the Calvin-Benson-Basham cycle are redox-regulated via the thioredoxin (Trx) system (Buchanan, 2016). In the chloroplast Trxs are reduced in the light by ferredoxin in a reaction catalyzed by the Ferredoxin Thioredoxin Reductase FTR. NTRC, a protein with both a NADPH-thioredoxin reductase and a Trx domain, has also been found in chloroplasts (Serrato et al., 2004). NTRC has been shown to be involved in controlling the level of H_2_O_2_ via its interaction with 2-cys peroxiredoxin (2-Cys-PRX), it participates in redox regulation of a key enzyme in starch biosynthesis (Michalska et al., 2009; Lepistö et al., 2013), of enzymes of the chlorophyll biosynthesis pathway (Richter et al., 2013; Perez-Ruiz et al., 2014) and in regulating the activity of the chloroplast ATP-synthase (Naranjo et al., 2016; Carrillo et al., 2016; Nikkanen et al., 2016; for a recent review see Cejudo et al., 2019). A severe growth inhibition phenotype has been reported for a mutant lacking NTRC and Trx f1 (Thormählen et al., 2015) and in the triple mutant *ntrc-trxf1f2* and the double mutant *ntrc-trxx* (Ojeda et al., 2017a). Both, *ntrc-trxf1f2* and *ntrc-trxx* mutants showed a high mortality at the seedling stage (Ojeda et al., 2017b) indicating that NTRC is important for chloroplast redox regulation, for controlling the redox state of several thioredoxins and showing that both redox regulation systems, FTR and the NTRC, are linked via 2-Cys-Prxs (Perez-Ruiz et al., 2017). Interaction between NTRC and Trx m1, Trx m3 and Trx y1 have been shown by bifluorescence complementation assays in vivo (Nikkanen et al., 2016). Trx m4 has been suggested to regulate negatively cyclic electron flow around photosystem I (PSI) (Courteille et al., 2013). Recently, evidence has been provided that m-type thioredoxins form a complex with PGRL1, a protein that is supposed to participate together with PGR5 in cyclic electron flow (Okegawa and Motohasi, 2020; Wolf et al., 2020). Trx m1, Trx m2 and Trx m4 have also been reported to be implicated in the biogenesis of PSII (Wang et al., 2013), showing a broad implication of m-type thioredoxins in the photosynthetic light reactions. Compared to the other isoforms, Trx m3 seems not to be relevant for controlling these processes. A mutant of Trx m3, the less abundant protein of the m-type thioredoxins in chloroplast stroma (Okegawa and Motohasi, 2015), shows unaltered chloroplast performance (Benítez-Alfonso et al., 2009). Tobacco plants overexpressing Trx m were shown to be impaired in photosynthesis but being more resistant to oxidative stress conditions (Rey et al., 2013). Furthermore, Trx m1, Trx m2 and NTRC have been shown to be indispensable for acclimation of photosynthesis under fluctuating light conditions (Thormählen et al., 2017). Taken together, these reports provide experimental evidence for an important role played by NTRC and Trx m isoforms for the rapid acclimation of plants to changes in the light regime, for controlling alternative photosynthetic electron flow and for coping with oxidative stress. Another important protein in the thiol-based chloroplast redox regulatory network is 2-Cys peroxiredoxin (2-Cys PRX), the most abundant peroxiredoxin in the chloroplast (Muthuramalingam et al., 2009). 2-Cys PRXs are reduced by NTRC and by Trxs and detoxify H_2_O_2_. Most importantly, they also act as a system to reoxidize reduced thioredoxins (Telman et al., 2020).

In the thylakoid lumen, candidates for controlling the redox state of protein disulphide bridges are the Lumen Thiol Oxidoreductase 1 (LTO1) and the atypical cytochrome *c*_6*A*_ (Marceida et al., 2006). The LTO1 protein is a transmembrane protein carrying a C-terminal thioredoxin-like domain typical of oxidoreductases belonging to the protein disulphide isomerase family. The *lto1* mutant had been shown previously to be affected in the assembly of active PSII while PSI electron transport was unaltered upon excitation with far-red light (Karamoko et al., 2011). Since the cysteine residues of LTO1 are at the lumen side of the thylakoid membrane, the redox state of the stroma has to be transmitted to these cysteines. Possible candidates are the transmembrane proteins CCDA and HCF 164 (Karamoko et al., 2013; Kang and Wang, 2016; Motohasi and Hisabori, 2010).

There is a strong link between the level of reactive oxygen species (ROS) and the thiol system. Superoxide anion radicals (O_2_^•-^) are mainly generated by the photosynthetic electron transport at photosystem I (PSI) by the classical “Mehler reaction” or pseudocyclic electron flow. It has been reported that leaves of *A. thaliana* and *N. tabacum* plants grown under short day conditions (SD, 8 h light, 16 h dark) have the double amount of superoxide compared with plants grown under long day conditions (LD, 16 h light, 8 h dark) (Michelet and Krieger-Liszkay, 2012). This extra electron transport in SD plants is used to generate a higher proton gradient and more ATP than found in thylakoids from LD plants. In the presence of an uncoupler, the difference in ROS generation was abolished between the two different thylakoid preparations. Addition of NADPH but not of NADH increased the level of ROS generation in LD thylakoids to the same amount as observed in SD thylakoids. Addition of NADPH to SD thylakoids had no significant effect. In thylakoids from plants lacking NTRC the ROS production was like in SD wild type (wt) thylakoids and the difference between SD and LD thylakoids was abolished (Lepistö et al., 2013). These results point to a redox regulation of ROS generation at the level of PSI.

It remains an open question whether NTRC interacts with a protein of PSI at the thylakoid membrane, whether a Trx is involved in the redox regulation of ROS generation at PSI and how the redox state of the stroma is transmitted to the thylakoid lumen. In this study we aimed to establish the interaction between different players of the chloroplast thioredoxin network and their ability to alter the capacity of O_2_ reduction at the PSI acceptor side. We measured light-dependent ROS generation on leaves and isolated thylakoids of *A. thaliana* grown under SD or LD conditions in wt, in single mutants: *ntrc, trxm4, ccda, lto1, cyt c*_*6A*_, in double mutants: *trxm1trxm2* and *2cpab*, and in plants overexpressing NTRC (oeNTRC). The sub-chloroplast localization of the NTRC and Trx m was studied using immunoblots. In vitro reconstitution experiments were performed using thylakoids and purified recombinant Trx m and/or NTRC in order to directly test their effect on light-induced ROS generation.

## Materials and Methods

### Plant Material

*A. thaliana* wt (Col-0) and mutants were grown for 6 weeks in soil either under short day conditions (8 h continuous white light - 160 μmol quanta m^-2^s^-1^, 21°C/16 h dark, 18°C) or long day conditions (16 h continuous white light - 160 μmol quanta m^-2^s^-1^, 21°C /8 h dark, 18°C). All mutants and over-expressing plants used were already described in previous studies: the *trxm4* T-DNA mutant (Courteille et al., 2007) and the *trxm1m2* mutant (Thormählen et al., 2017); the T-DNA insertion mutant of NTRC (*ntrc*) (Lepistö et al., 2009) and transgenic plants overexpressing wild type NTRC protein in *ntrc* background (Toivola et al., 2013); the double T-DNA mutant lacking the two 2-Cys Prxs, A and B (*2cpab*) (Ojeda et al., 2018); the T-DNA mutants lacking CCDA isoforms (Page et al., 2004); the T-DNA mutant lacking Cytcochrome *c*_*6A*_ (Pesaresi et al., 2009).

### Extraction of proteins from leaves

*Arabidopsis* shoots were grinded in liquid nitrogen before homogenization in lysis buffer. The lysis buffer contained 100 mM Tris–HCl pH 6.8, 4% sodium dodecyl sulphate (SDS), 20 mM EDTA and protease inhibitor cocktail (Sigma-Aldrich, St. Louis, MI, USA).

### Extraction of thylakoids from A. thaliana

Young fully expanded leaves were grinded in 0.33 M sorbitol, 50 mM KCl, 10 mM EDTA, 1 mM MgCl_2_, 25 mM Mes pH 6.1. After centrifugation, the pellet was washed twice with 0.33 M sorbitol, 60 mM KCl, 2 mM EDTA, 1 mM MgCl_2_, 25 mM HEPES pH 6.7. After centrifugation, the pellet was resuspended in 0.3 M sucrose, 50 mM KCl, 1 mM MgCl_2_, 20 mM HEPES pH 7.6 (measurement buffer). This procedure was repeated once, and the pellet was resuspended to a final concentration of about 1 mg of chlorophyll per ml of thylakoids. All centrifugations were performed at 3,000xg for 3 min at 4°C.

### Isolation of Photosystem I

Photosystem I was isolated as described in Krieger-Liszkay et al. (2020).

### Room-Temperature Spin-Trapping EPR Measurements

Spin-trapping assays with 4-pyridyl-1-oxide-*N*-*tert*-butylnitrone (4-POBN) (Sigma-Aldrich) were carried out using leaf disks or freshly isolated thylakoid membranes at a concentration of 10 μg of Chl ml^-1^. Leaf disks were vacuum-infiltrated with the buffer containing the spin trap reagents prior to the illumination and then floating on the same buffer during the illumination. Samples were illuminated for a given time with white light (200 μmol quanta m^-2^ s^-1^ in case of leaf disks and 500 μmol quanta m^-2^ s^-1^ in case of thylakoids) in the presence of 50 mM 4-POBN, 4% ethanol, 50 μM Fe-EDTA, and buffer (25 mM HEPES, pH 7.5, 5 mM MgCl_2_, 0.3 M sorbitol). When indicated, 200 μM NADPH, 0.3 μM Trxm4 and 0.3 μM NTRC were added to the assay before starting the illumination.

EPR spectra were recorded at room temperature in a standard quartz flat cell using an ESP-300 X-band (9.73 GHz) spectrometer (Bruker, Rheinstetten, Germany). The following parameters were used: microwave frequency 9.73 GHz, modulation frequency 100 kHz, modulation amplitude: 1G, microwave power: 6.3 milliwatt in 4-POBN assays, receiver gain: 2x10^4^, time constant: 40.96 ms; number of scans: 4.

### O_2_ consumption

Measurements of O_2_-consumption were performed in a Liquid-Phase Oxygen Electrode Chamber (Hansatech Instruments, Norfolk, England) using isolated PSI (10 μg Chl ml^−1^) in Tricine 20 mM pH 8.0, in the presence of 5 mM MgCl_2_, 30 mM NaCl and 5 mM ascorbate, 30 μM 2,6-dichlorophenolindophenol (DCPIP) as exogenous electron donors to P700^+^.

### SDS-PAGE and Western Blotting

SDS-PAGE was performed using 8% or 4-20 % polyacrylamide gels. Proteins were blotted onto a nitrocellulose or a PVDF membrane. Labelling of the membranes with polyclonal antibodies, produced in the lab (anti-Trxm; anti-NTRC; anti-2-Cys PRX) or commercially available (PsaF and β- subunit of ATP synthase; Agrisera, Vännäs, Sweden), was carried out at room temperature in 50 mM Tris–HCl pH 7.6, 150 mM NaCl, 0.1% Tween-20 and 5% non-fat milk powder. After washing, bound antibodies were revealed with a peroxidase-linked secondary anti-rabbit antibody (Agrisera, Vännäs, Sweden) and visualized by enhanced chemiluminescence.

### Redox state of PsaF

Total leaf protein samples were prepared as described above with AMS or mPEG-maleimide (2 mM) added to the extraction buffer. 50 μg protein samples were electrophoresed (4-20 % SDS-PAGE) and PsaF was immuno-detected (PVDF membrane / Chemiluminescence) and signals corresponding to reduced (alkylated) and oxidized forms were quantified using ImageLab software (BioRad).

### NTRC Trx reduction assays

TRX reduction tests by NTRC were performed using DTNB as already described in Bohrer et al. (2012).

## Results

Leaves and thylakoid membranes isolated from *A. thaliana* wild-type (wt) plants grown in short day (SD) conditions generate in the light about twice the amount of ROS compared to those from plants grown in long day (LD) conditions (Fig. 1) as has been previously shown (Michelet and Krieger-Liszkay, 2012). ROS production was measured using an indirect spin trapping assay. In this assay hydroxyl radicals are detected which derive from superoxide anion radicals and hydrogen peroxide in a Haber-Weiss reaction catalyzed by Fe(II) (Michelet and Krieger-Liszkay, 2012). A comparison between ROS production in leaf disks and isolated thylakoids shows that the same differences in ROS production are found in both types of samples (Fig. 1B, C). This demonstrates that in the light the majority of superoxide/hydrogen peroxide is generated by the photosynthetic electron transport chain. Mutants affected in Trx m isoforms or in NTRC lost the difference between SD and LD. The single mutant *trxm4* and the double mutant *trxm1m2* generated similar amounts of ROS like LD plants, while *ntrc* generated high amounts of ROS like SD plants, independently of the photoperiod during their growth. Overexpression of NTRC increased the difference between SD and LD compared to wt. In a similar manner the mutant devoid of 2-Cys PRX A and B showed an overall increase in the ROS production, however, the difference between SD and LD was maintained in *2cpab* (Fig. 1).

**Figure 1.**
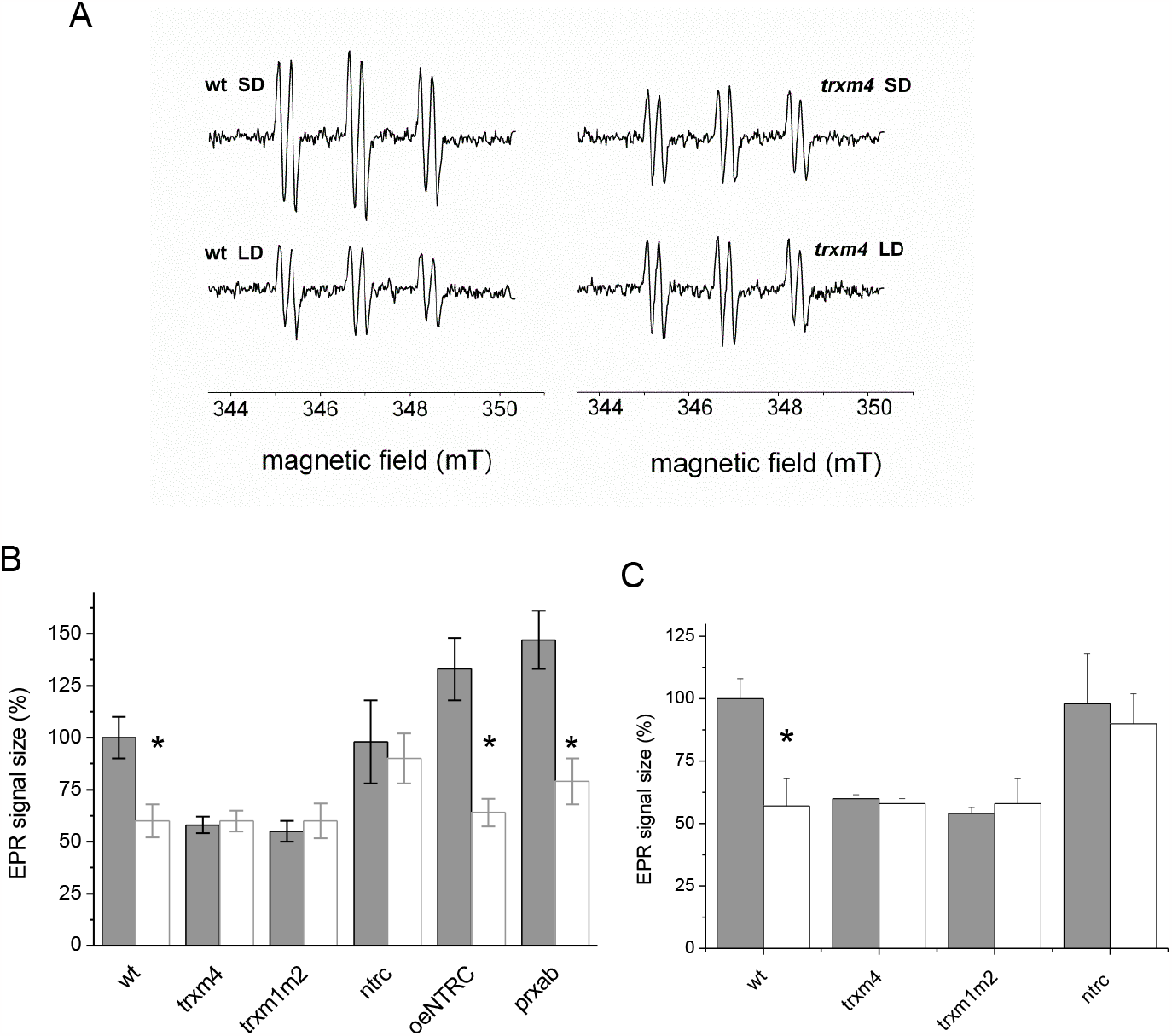
Light-induced hydroxyl radical formation in SD and LD leaf disks and washed thylakoid membranes in wild type and redoxin mutants. Generation of hydroxyl radicals originating from O_2_^•-^/H_2_O_2_ was detected by spin trapping with 4-POBN. A: Typical EPR spectra of the 4-POBN/α-hydroxyethyl adduct are shown. After infiltration with assay medium containing 4-POBN/ethanol/FeEDTA, leaf disks were incubated in the same medium for 30 min in light (200 μmol quanta m^-2^s^-1^) before detection of the radicals in the medium. Difference in EPR signal size for leaf samples (B) and thylakoid membranes (C). Thylakoid membranes (20 μg ml^-1^) were illuminated for 2 min (500 μmol quanta m^-2^s^-1^) in the presence of the spin trapping assay before detection of the radical. Grey bars: SD, white bars: LD. All EPR signals were normalized to the signal of SD thylakoids without addition (100%). Mean values are shown (n≥6, biological replicates; *, P <0.05 (comparison between SD and LD growth conditions for each genotype) according to Tukey test.

Furthermore, we found that 2-Cys PRX was slightly more abundant in leaf protein samples from plants grown in LD than in SD when prepared in presence of SDS but not in absence of detergent (Fig. 2A and B), suggesting that 2-Cys PRX is mainly stromal and the difference between SD and LD samples would be attributable to a differential association to thylakoid membranes. Indeed, thylakoids from LD grown plants showed a marked higher amount of 2-Cys PRX (recovered from thylakoid membranes in the presence of SDS) compared to SD, confirming that the difference of global (stromal plus thylakoid-associated) abundance in leaf extracts between photoperiods was attributable to the fraction of membrane bond protein. In LD, the protein was present mainly in its dimeric form, most probably corresponding to the oxidized form as evidenced by the shift of the signal from an apparent mass of about 40 kDa to about 20 kDa reproduced by reduction with DTT giving the monomeric form. In PSI preparations, we could not immune-detect 2-Cys PRX (Fig. 2C) suggesting that the protein was not directly associated to PSI complexes, or lost upon sample preparation due to a loosen association to the photosystem. This latter possibility is strongly suggested by the detection of a faint signal in the supernatant of LD thylakoids resuspended in buffer devoid of detergent.

**Figure 2.**
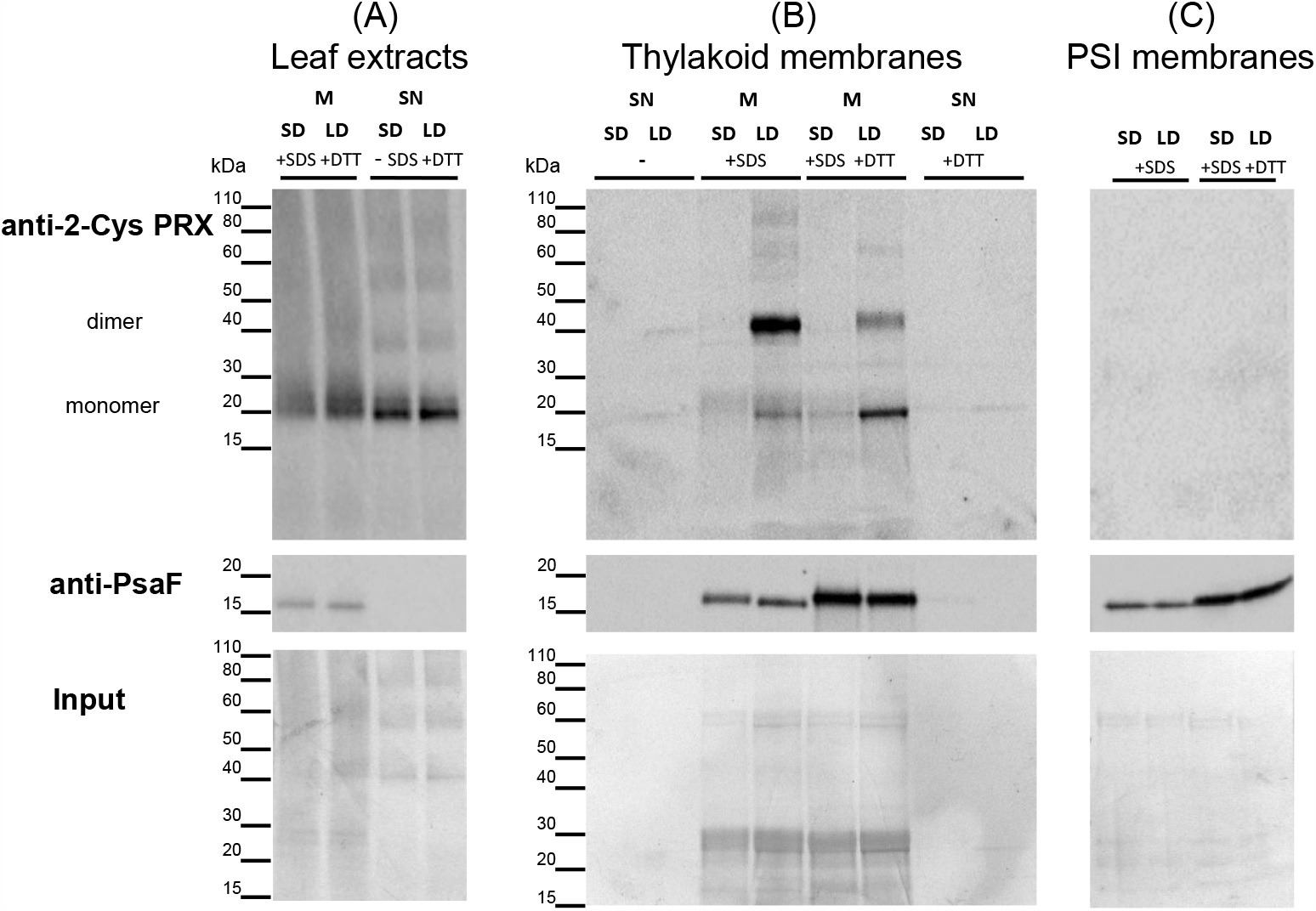
Association of 2-Cys PRX with thylakoid membranes depends on the photoperiod. Immuno-detection of 2-Cys PRX in (A) leaf extracts (50 μg protein per well), (B) thylakoid membranes (5 μg chl per well) or (C) PSI complexes (0.5 μg chl per well) prepared from plants grown either in short day (SD) or long day (LD) conditions. Immuno-detection of PsaF and coomassie staining (input) were taken as membrane-associated protein extraction and gel loading controls, respectively. SDS (2%) and DTT (10 mM) treatments of membrane samples (M) were performed for 10 min at RT. After centrifugation the supernatant (SN) was collected and supplemented with non-reducing gel loading blue prior heat treatment and gel loading.

To see whether the amount of Trx m and/or NTRC is altered in plants grown in the two different light regimes, immunoblots were performed using leaf extracts and isolated thylakoids. As shown in Fig. 3, no difference in the total amount of Trx m4 and NTRC was found in leaf extracts. However, the attachment of Trx m differed in SD and LD thylakoids. Trx m4 was found in the thylakoid fraction in SD thylakoids but not in LD thylakoids. A similar membrane localization like for Trx m4 was observed for Trx m2 (Sup. Fig. 1).

**Figure 3.**
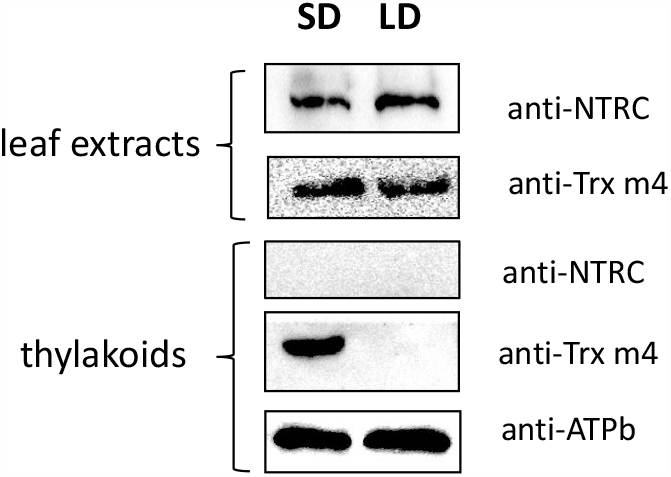
Membrane association of Trx m4 to thylakoid membranes in SD conditions. Immuno-detection of NTRC and Trx m4 in leaf extracts and thylakoid membranes prepared from plants grown either in short day (SD) or long day (LD) conditions. Samples corresponding to 5 μg chlorophyll were loaded in each well. The results shown are representative of three biologically independent experiments.

Attachment of Trx m seems to be required for allowing electron transport to oxygen at PSI. To test whether the lack of membrane-associated Trx was indeed responsible for the lower ROS generation in LD thylakoids, we reconstituted LD thylakoids with purified Trx m4 and/or NTRC proteins in the presence of NADPH (Fig. 4). Addition of NADPH alone stimulated slightly the ROS production in LD thylakoids. A further increase in signal size was observed when Trxm4 or NTRC were added together with NADPH. However, these differences were statistically not significant. When Trxm4 was added together with NADPH and NTRC, LD thylakoids generated three-fold more ROS than without any protein addition. In the absence of NADPH, addition of TRxm4 and NTRC had no effect. The different additives had only a small effect on SD thylakoids. This result points to a redox regulation of O_2_ reduction at the level of PSI. To investigate whether there is a direct effect of thiol reduction on the O_2_ reduction capacity of PSI, we incubated isolated PSI with the reducing agent TCEP and followed light-dependent O_2_ consumption using an O_2_ electrode. As shown in Fig. 5, O_2_ consumption was two times higher in the presence of TCEP. These data show that it is redox regulation of PSI itself that is crucial for the level of O_2_ reduction at the acceptor side of PSI.

**Figure 4.**
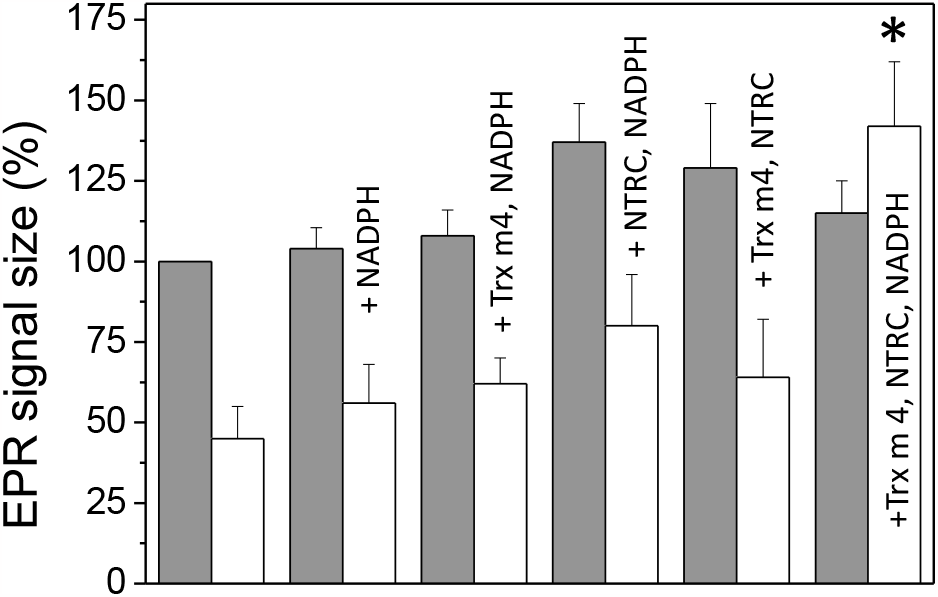
Reconstitution of ROS generation in LD thylakoids by Trxm, NTRC and NADPH. Light-induced hydroxyl radical formation in SD and LD thylakoids is shown by indirect spin trapping with 4-POBN. Thylakoid membranes (20 μg ml^-1^) were illuminated for 2 min (500 μmol quanta m^-2^s^-1^) before detection of the radical. When indicated, 200 μM NADPH, 0.3 μM Trx m4 and 0.3 μM NTRC were added to the assay before starting the illumination. All EPR signals were normalized to the signal of SD thylakoids without addition (100%). Grey bars: SD, white bars: LD. Mean values are shown (n=3; *, P <0.05 (comparison with LD no protein added) according to Tukey test.

**Figure 5.**
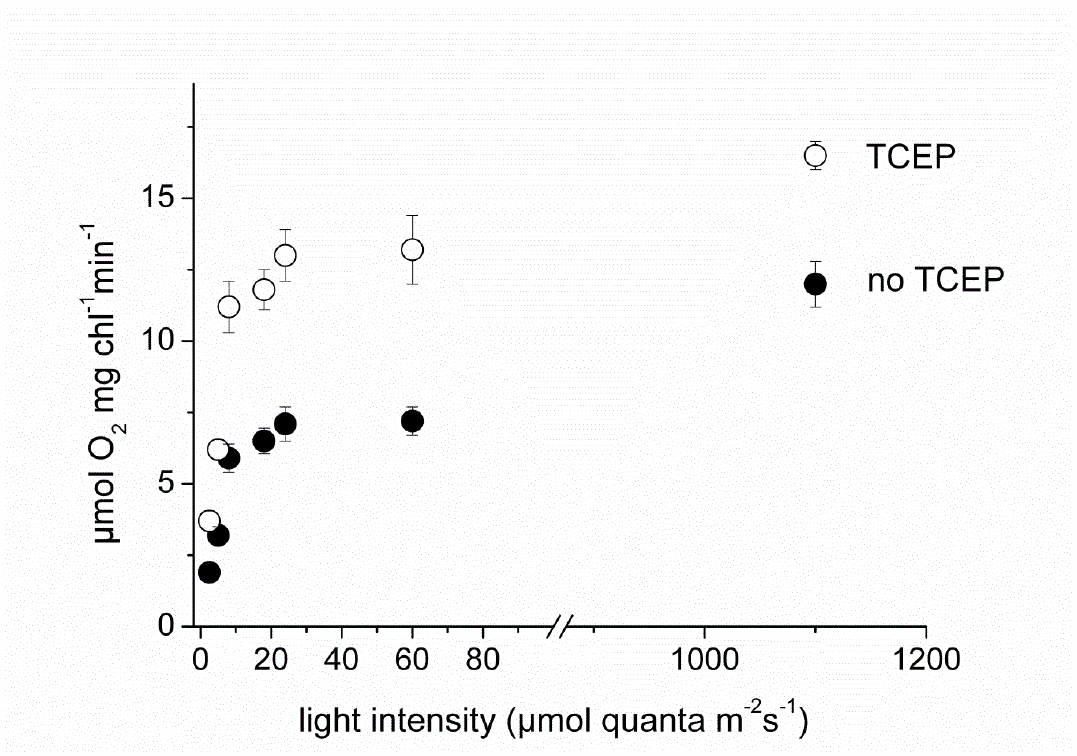
O_2_ consumption by isolated PSI as a function of the light intensity. O_2_ consumption was measured with an O_2_ electrode using DCPIP/ascorbate as electron donor. Samples were illuminated with green actinic light. When indicated, isolated PSI had been incubated with 1 mM TCEP for 15 min prior to the measurement.

The question arises which subunit of PSI may be redox-regulated. Several subunits of PSI contain cysteines, however most of them are ligands of the iron-sulphur clusters F_x_, F_A_ and F_B_ and can therefore be excluded as candidates undergoing reversible regulatory redox modifications. In addition, there are two proteins, PsaN and PsaF that may be candidates for disulfide bridge formation. It has been shown previously that the PSI subunit PsaN contains four cysteine residues that can form two disulfide bridges (Motohashi and Hisabori, 2006). PsaF contains at its luminal site 2 cysteine residues that are close enough to form a disulfide bridge. PsaF is a transmembrane protein that forms the docking site for plastocyanin at the donor side of PSI and at the acceptor side its C-terminus is in direct neighborhood with PsaE. PsaE forms together with PsaC and PsaD the docking site for ferredoxin. O_2_ reduction is supposed to take place at this site. Since PsaF is the most likely candidate for redox regulation, and we did not observe any significant quantitative difference of this protein when comparing SD and LD thylakoid and PSI protein preparations (Fig. 2), we performed redox western assays to explore the influence of light regime on its redox state. Thylakoid membranes were treated with 4′-acetamido-4′-maleimidylstilbene-2,2′-disulfonic acid (AMS) that reacts with thiol groups (alkylation reaction), we separated the proteins by SDS-PAGE and analysed the apparent mass of PsaF by immunodetection. Sup. Fig. 2 shows that two bands were detected when the thylakoids had been chemically reduced with tris(2-carboxyethyl) phosphine prior to the AMS treatment. This indicates that PsaF exists in an oxidized form with cysteines potentially forming a disulfide bridge and, upon addition of a reductant, in a reduced form with AMS accessible thiols. However, without exogenous reductant, in isolated thylakoids from both SD or LD plants only the oxidized form was found, probably due to spontaneous oxidation during membrane sample preparation. Redox Westerns were performed on the mutants used for detection of the ROS levels in Fig. 1. In leaf extracts prepared in presence of SDS and m-PEG-mal (Fig. 6, Sup. Fig. 3) PsaF was immuno-detected as two well distinguishable redox variants that could be quantified by densitometry. We found that relative abundance of PsaF oxidized and reduced forms strongly varied between light and dark samples. In LD, PsaF was reduced equally in all genotypes with a reduction percentage ranging from 24.6 to 32.4 % in the dark, and from 1.5 to 2.9 % in the light. But, we did not observe significant differences in the redox state of PsaF in the two light regimes or between the different mutant lines, with the exception of *ntrc* where PsaF was significantly more reduced in SD (39.7 % in the dark and 8.4 % in the light) than in LD (24.6 % in the dark and 2.4 % in the light), and in comparison to wt (28.4 % in the dark and 2.2 % in the light) and the other mutants analyzed. However, there was no correlation with the ROS levels found in *ntrc*. We concluded that the reduction of PsaF may be not stable enough to catch it during the alkylation treatment procedure.

**Figure 6.**
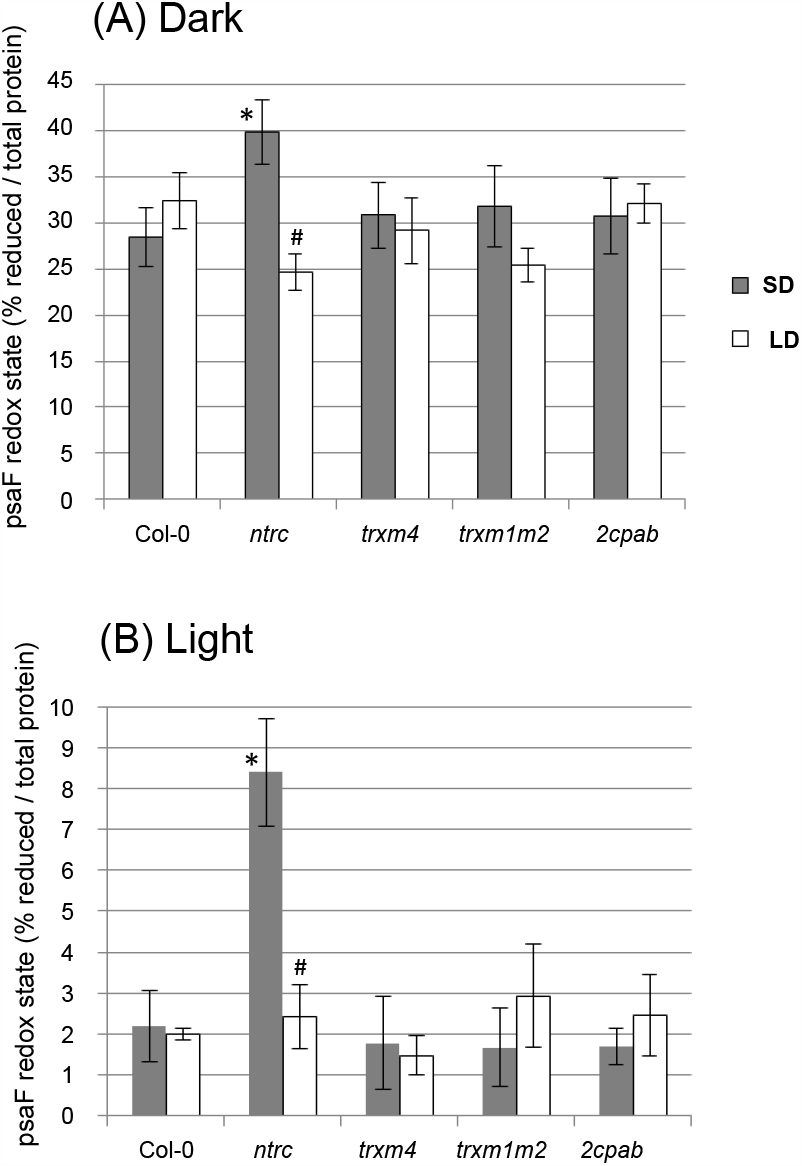
Redox state of PsaF in dark- and light-adapted plants. Total leaf protein extracts from WT (Col-0) and redoxins (*ntrc, trxm4, trxm1m2* and *2cpab*) mutant plants grown in SD or LD conditions were prepared in presence of the thiol alkylating agent mPEG-maleimide. After SDS-PAGE, proteins (50 μg per well) were electro-transferred onto a PVDF membrane for PsaF immuno-detection (chemi-luminescence). See Sup. Fig. 3 for details about the quantification method used and an example of redox western signals. (A) and (B) PsaF redox state in leaves in the dark and in the light, respectively. Values correspond to the mean of 5-6 experiments. Error bars correspond to standard deviation. * and # designate significant differences between mutant and wt genotypes, and between SD and LD (for each genotype), respectively; according to Student’s t-test, P < 0.05.

The redox state of the stroma has to be transmitted to the thylakoid lumen to be able to act on thiol/disulphide groups of PsaF. The transmembrane proteins CCDA and HCF164 are likely candidates for the transmission of the redox state from the stromal site to the lumenal site of the thylakoid membrane (Motohashi and Hisabori, 2010). LTO1 (Karamoko et al., 2011) and the atypical cytochrome *c*_*6A*_ (Marceida et al., 2006) may act as candidates for redox modifications inside the lumen. Fig. 7 shows that the mutants of CCDA (*ccda3* and *ccda4*) and of LTO1 have indeed lost the difference in superoxide production between SD and LD, while loss of cyt *c*_*6A*_ has no effect.

**Figure 7.**
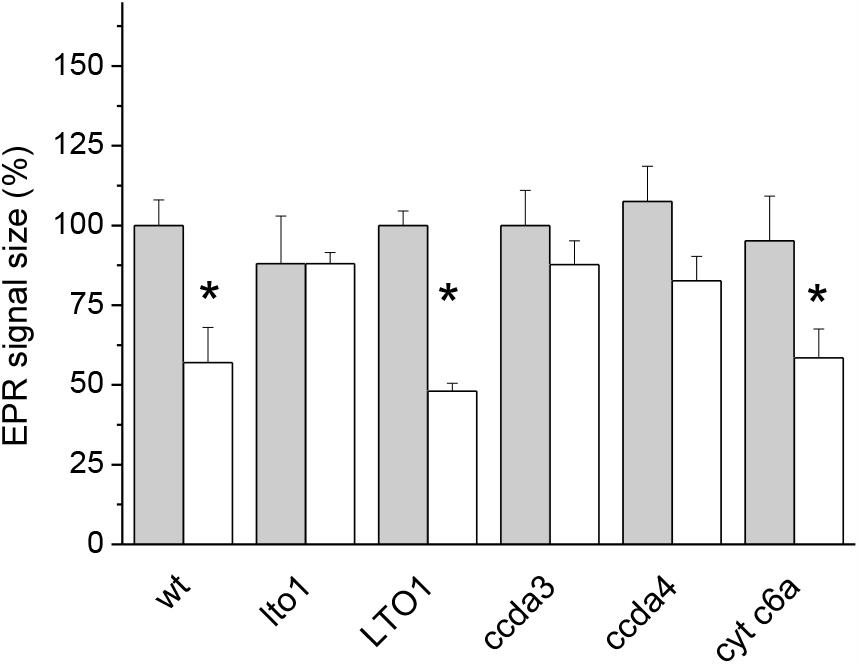
Light-induced hydroxyl radical formation in SD and LD leaf disks of mutants affected in LTO1, 2-Cys-PRX (*prxab*), CcdA (*ccda3* and *ccda4*) and Cyt *c*_6*A*_. Generation of hydroxyl radicals originating from O_2_^•-^/H_2_O_2_ was detected in leaves by spin trapping with 4-POBN. Grey bars: SD, white bars: LD. All EPR signals were normalized to the signal of SD thylakoids without addition (100%). Mean values are shown (n=8, 2-3 biological replicates; *, *P* <0.05 (comparison between growth conditions for each genotype) according to Tukey test.

## Discussion

Redox regulation of linear photosynthetic electron transport and alternative pathways has been reported previously (Johnson, 2003). Courteille and coworkers (2013) showed that cyclic electron transport involving the NDH complex was altered in Trx m4 mutants, Nikkanen et al. (2018) reported the involvement of NTRC in the control of NDH complex-dependent and Naranjo et al. (2021) in PGR5-dependent cyclic flow. Both alternative electron transport pathways, cyclic and pseudocyclic flow, are in competition and down-regulation of cyclic flow is therefore supposed to stimulate pseudocyclic flow. We have shown previously (Michelet and Krieger-Liszkay, 2012) that O_2_ reduction at PSI is higher in SD thylakoids than in LD thylakoids and that this difference is abolished in the presence of uncouplers. This observation points to a pH- or redox-regulated process taking place at the luminal side of the thylakoid membrane and controlling O_2_ reduction at PSI. Indeed, O_2_ reduction of isolated PSI is stimulated by the thiol-reducing agent TCEP (Fig. 5). At low light intensities, such as those used for the measurements shown in Fig. 5, O_2_ is reduced by the terminal electron acceptors, the 4Fe4S clusters F_A_ and F_B_, while at higher light intensities it is reduced by the acceptor A_1_, a phylloquinone (Kozuleva et al., 2021). PsaN and PsaF are the only constitutive protein subunits of PSI containing cysteine residues that can form a disulphide bridge. The most likely candidate for the redox modification affecting O_2_ reduction is PsaF for the following reasons: 1. PsaF was found in PSI x-ray structures from pea either in the reduced state (Mazor et al., 2015) or with a disulfide bond (Qin et al., 2015), and 2. PsaF has been identified by a proteomics approach as a redox-affected protein (Ströher and Dietz, 2008). PsaF is a transmembrane protein, and we hypothesize here that, upon a modification of the redox state of cysteines, a long-range structural change affects the neighboring subunit PsaE and thereby the ferredoxin docking site at the PSI acceptor side. Accordingly, O_2_ reduction is favored when the disulfide bridge in PsaF is reduced while Fd reduction becomes less efficient. The subunit PsaE seems to be crucial for O_2_ reduction at PSI in higher plants (Krieger-Liszkay et al., 2020). Unfortunately, we were not able to detect significant differences in the reduction level of PsaF in the two light regimes and in the mutant lines analyzed where distinct O_2_ reduction levels were clearly found (Fig. 1). We cannot rule out at present that PsaN may also be involved in the redox-dependent regulation of O_2_ reduction at PSI. Different to PsaF, PsaN is a peripheral luminal protein belonging to the light harvesting complex of PSI with no connection to the acceptor side of PSI. However, in the structure published by Pan et al. (2018) for maize PSI, PsaN forms close contacts with the N-terminal extension of PsaF. Redox modification of PsaN may impose a structural change on PsaF that then, as described above, may exert a structural alteration of the PSI subunits forming the Fd docking site.

We also addressed the question how the redox state of PsaF and/or PsaN is controlled in the lumen. *Lto1* mutant has lost the difference between SD and LD (Fig. 7), implying a role of this protein in redox regulation of PSI. The difference between SD and LD is also lost in the mutants *ccda3* and *ccda4* (Fig. 7), demonstrating the importance for CCDA in transmitting the stromal redox state to the lumen in vivo. Mutants of CCDA and LTO1 show high ROS levels independent of the growth photoperiod. We hypothesize that LTO1 keeps PsaF oxidized under LD conditions, while CCDA keeps LTO1 oxidized, leading to low O_2_ reduction in LD. In SD conditions, where the redox state of CCDA and LTO1 is highly reduced, the situation changes. Under these circumstances, reduction of PsaF (or/and PsaN) seems feasible (Fig. 8).

**Figure 8.**
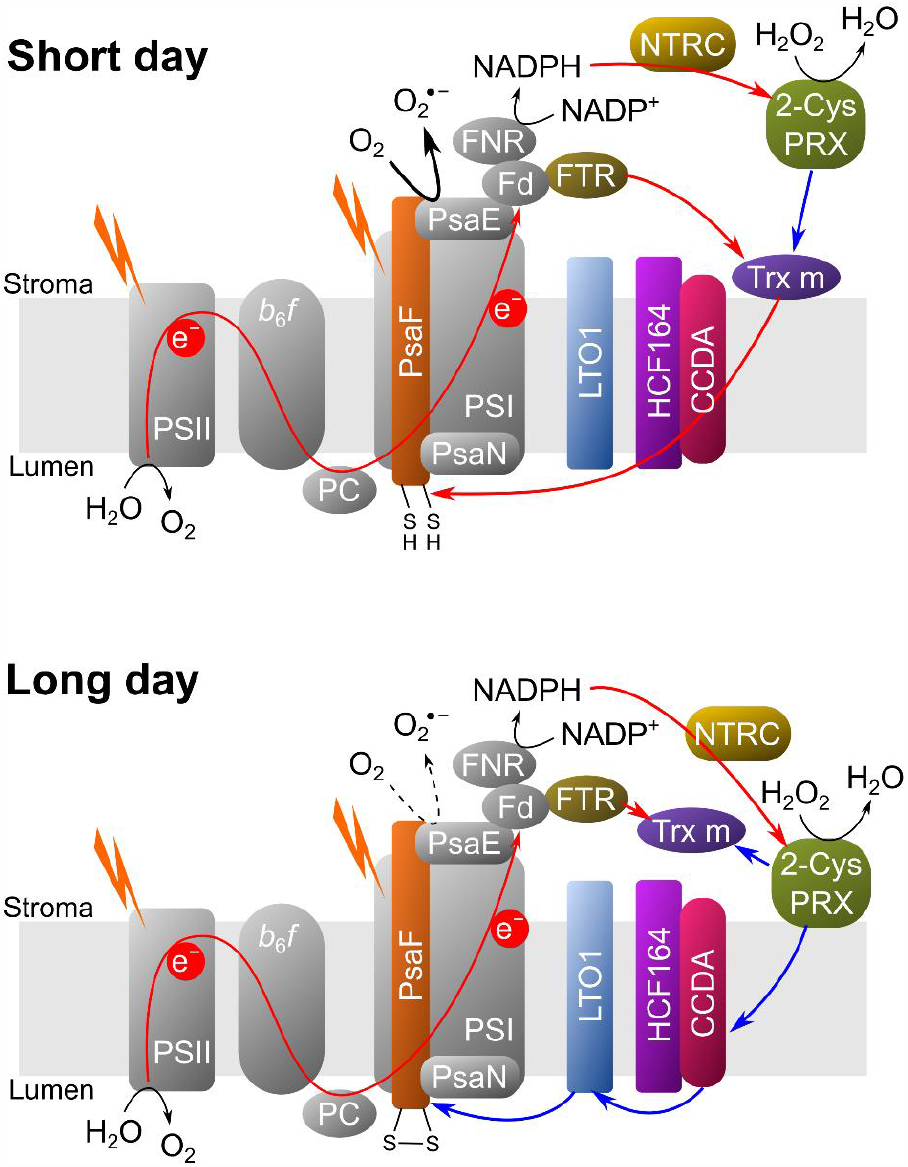
Model of redox regulation of O_2_ reduction at photosystem I. Reduced cysteine residues in PsaF favor higher O_2_ reduction activity. The redox state of PsaF is modified by two thiol modulating redox systems. The first one is required for reduction of the disulfide bond, the second one for their oxidation. Trx m is a central player for controlling the reduction state of PsaF. Trx m is reduced by FTR. Under SD conditions, Trx m associates to the thylakoid membrane and reduces CCDA which reduces via HCF164 finally PsaF. In LD conditions, reduced PsaF is oxidized by LTO1, which itself is oxidized by CCDA (blue arrows. 2-Cys-PRX attaches to the membrane, allowing the oxidation of CCDA. In addition, it can oxidize Trx m (blue arrows).

Trx m has been shown to be the electron donor to CCDA in vitro (Motohashi and Hisabori, 2010). We propose here that Trx m association to the thylakoid membrane is required for an efficient electron donation to CCDA and onwards to redox regulated proteins in the thylakoid lumen, reduction of PsaF (and/or PsaN) and increase in O_2_ reduction (Fig. 7). Trx m association to the membrane in SD conditions (Fig. 3 and Sup. Fig. 1) may be facilitated by the higher proton motive force generated in thylakoids from SD plants (Michelet and Krieger-Liszkay, 2012). Such a mechanism has been shown to play a role for the attachment of the plastid terminal oxidase to the thylakoid membrane (Bolte et al., 2020).

The question arises how to integrate 2-Cys-PRX and NTRC into this model. 2-Cys-PRXs are mainly reduced by NTRC and act as an oxidizing system towards reduced Trxs (Nikkanen et al., 2016; Perez-Ruiz et al., 2017; Telman et al., 2020). We suggest that membrane association of 2-Cys-PRX in LD is important to keep CCDA oxidized. As shown in Fig. 2, 2-Cys PRX is associated to the membrane only in LD conditions and is found mostly in its dimeric form. Besides controlling the redox state of Trxs, 2-Cys-PRX is responsible for H_2_O_2_ detoxification in the stroma (König et al., 2002). H_2_O_2_ detoxification may be more efficient if 2-Cys PRX is attached to the membrane close to the site of superoxide production and conversion into H_2_O_2_ by membrane-associated SOD. In the absence of NTRC, 2-Cys PRX is not able to fulfill this role, resulting in higher ROS levels independent of the photoperiod (Fig. 1). In the absence of 2-Cys PRX, as expected, the ROS levels are increased, but the difference between SD and LD is maintained (Fig. 1B). According to a previous report (Bohrer et al., 2012) and our measurements of DTNB reduction (Sup. Fig. 4), NTRC is not able to reduce Trxs m in a direct manner when tested at physiologically relevant concentrations.

In conclusion, our hypothesis for the redox regulation of pseudocyclic electron flow is based on three mechanisms: 1. The availability of electron acceptors other than O_2_ for photosynthetic electron transport, 2. Redox regulation of the different players according to the redox state of the stroma and the capabilities of the different players to form a network of redox-driven interactions and 3. The reversible membrane attachment of Trx m and 2-Cys-PRX that may depend on changes in pH and ion concentration controlled by the proton motive force. Attachment of Trx m4 seems to be necessary to achieve a high reduction state of CCDA and PsaF in SD resulting in high O_2_-reduction levels at PSI, while detachment of Trx m and attachment of 2-Cys-PRX in LD seems to favor oxidation of CCDA, and PsaF that is oxidized by LTO1, resulting in low levels of O_2_-reduction. However, the connection of redox regulation of PsaF *in vivo* with the photoperiod remains to be demonstrated.

Reduction of O_2_ in pseudocyclic electron flow is in competition with cyclic electron flow. Both pathways lead to the formation of a proton gradient without generating NADPH. According to Takahashi et al (2013), the redox state of the chloroplast controls the formation of supercomplexes composed of PSI, Cyt *b*_6_*f*, LHCs, PGRL1, FNR. Such supercomplexes are thought to be required for cyclic electron flow (Iwai et al., 2010). The formation of a complex between reduced Trx m and PGRL1 may inhibit cyclic electron flow by preventing the supercomplex formation required for cyclic flow. This suggestion is supported by the recent reports on the interaction between Trx m and PGRL1 (Wolf et al., 2020; Okegawa and Motohashi, 2020). As shown recently, Trx x and Trx y play an important role in in the acceptor-side regulation of PSI and protection of PSI against photoinhibition under fluctuating light conditions (Okegawa et al., 2023). Future work on the extent of cyclic flow and superoxide production in mutants of the different components of the Trx system will show if both, pseudocyclic and cyclic flow are controlled by the same proteins but in the opposite way.

## Acknowledgements

We would like to thank Patrice Hamel (Ohio State University, USA) for sending us the seeds of the *lto1* mutants. This work was supported by the Labex Saclay Plant Sciences-SPS (ANR-17-EUR-0007) and the platform of Biophysics of the I2BC supported by the French Infrastructure for Integrated Structural Biology (FRISBI; grant number ANR-10-INSB-05). U.H. is supported by a CNRS PhD fellowship.

## Author Contribution

E.I-B. and A.K-L. designed the project. U.H., B.N., G.S., C.E., H.V., E.I-B. and A.K-L. performed the experiments and analysed the data. P.S. and E.R. participated in discussions. A.K-L. wrote the initial version of the manuscript that was read and revised by all authors.

## Data availability

Data will be made available on demand.

## References

Benitez-Alfonso Y, Cilia M, San RA, Thomas C, Maule A, Hearn S, Jackson D (2009) Control of Arabidopsis meristem development by thioredoxin-dependent regulation of intercellular transport. Proc. Natl. Acad. Sci. U. S. A. 106: 3615–3620.

Bohrer AS, Massot V, Innocenti G, Reichheld JP, Issakidis-Bourguet E, Vanacker H (2012) New insights into the reduction systems of plastidial thioredoxins point out the unique properties of thioredoxin z from Arabidopsis. J Exp Bot 63: 6315–23.

Bolte S, Marcon E, Jaunario M, Moyet L, Paternostre M, Kuntz M, Krieger-Liszkay A (2020) Dynamics of the localization of the plastid terminal oxidase inside the chloroplast. J Exp Bot. 71: 2661–2669.

Buchanan BB (2016) The path to thioredoxin and redox regulation in chloroplasts. Annu. Rev. Plant Biol. 67: 1–24.

Carrillo LR, Froehlich JE, Cruz JA, Savage LJ, Kramer DM (2016) Multi-level regulation of the chloroplast ATP synthase: the chloroplast NADPH thioredoxin reductase C (NTRC) is required for redox modulation specifically under low irradiance. Plant J. 87: 654–663.

Cejudo FJ, Ojeda V, Delgado-Requerey V, González M, Pérez-Ruiz JM (2019) Chloroplast Redox Regulatory Mechanisms in Plant Adaptation to Light and Darkness. Front Plant Sci. 10: 380.

Courteille A, Vesa S, Sanz-Barrio R, Cazale AC, Becuwe-Linka N, Farran I, Havaux M, Rey P, Rumeau D (2013) Thioredoxin m4 controls photosynthetic alternative electron pathways in Arabidopsis. Plant Physiol. 161: 508–520.

Iwai M, Takizawa K, Tokutsu R, Okamuro A, Takahashi Y, Minagawa J (2010) Isolation of the elusive supercomplex that drives cyclic electron flow in photosynthesis. Nature 464: 1210–3.

Johnson GN (2003) Thiol regulation of the thylakoid electron transport chain--a missing link in the regulation of photosynthesis? Biochemistry 42: 3040–4.

Kang ZH, Wang GX (2016) Redox regulation in the thylakoid lumen. J. Plant Physiol. 192: 28–37.

Karamoko M, Cline S, Redding K, Ruiz N, Hamel PP (2011) Lumen Thiol Oxidoreductase1, a disulfide bond-forming catalyst, is required for the assembly of photosystem II in Arabidopsis. Plant Cell 23: 4462–75.

Karamoko M, Gabilly ST, Hamel PP (2013) Operation of trans-thylakoidthiol-metabolizing pathways in photosynthesis. Front. Plant Sci. 4: 476.

König J, Baier M, Horling F, Kahmann U, Harris G, Schürmann P, Dietz KJ (2002) The plant-specific function of 2-Cys peroxiredoxin-mediated detoxification of peroxides in the redox-hierarchy of photosynthetic electron flux. Proc Natl Acad Sci U S A 99: 5738–43.

Krieger-Liszkay A, Shimakawa G, Sétif P (2020) Role of the two PsaE isoforms on O2 reduction at photosystem I in Arabidopsis thaliana. Biochim Biophys Acta Bioenerg. 1861: 148089

Lepistö A, Kangasjärvi S, Luomala EM, Brader G, Sipari N, Keränen M, Keinänen M, Rintamäki E (2009) Chloroplast NADPH-thioredoxin reductase interacts with photoperiodic development in Arabidopsis. Plant Physiol. 149: 1261–76.

Lepistö A, Pakula E, Toivola J, Krieger-Liszkay A, Vignols F, Rintamaki E (2013) Deletion of chloroplast NADPH-dependent thioredoxin reductase results in inability to regulate starch synthesis and causes stunted growth under short-day photoperiods. J. Exp. Bot. 64: 3843–3854.

Marcaida MJ, Schlarb-Ridley BG, Worrall JA, Wastl J, Evans TJ, Bendall DS, Luisi BF, Howe CJ (2006) Structure of cytochrome c6A, a novel dithio-cytochrome of Arabidopsis thaliana, and its reactivity with plastocyanin: implications for function. J Mol Biol 360: 968–77.

Mazor Y, Borovikova A, Nelson N (2015) The structure of plant photosystem I super-complex at 2.8 A resolution. Elife: 4:e07433. doi: 10.7554/eLife.07433

Michalska J, Zauber H, Buchanan BB, Cejudo FJ, Geigenberger P (2009) NTRC links built-in thioredoxin to light and sucrose in regulating starch synthesis in chloroplasts and amyloplasts. Proc. Natl. Acad. Sci. U. S. A. 106: 9908–9913.

Michelet L, Krieger-Liszkay A (2012) Reactive oxygen intermediates produced by photosynthetic electron transport are enhanced in short-day grown plants. Biochim Biophys Acta 1817: 1306–13.

Motohashi K, Hisabori T (2006) HCF164 receives reducing equivalents from stromal thioredoxin across the thylakoid membrane and mediates reduction of target proteins in the thylakoid lumen. J. Biol. Chem. 281: 35039–35047.

Motohashi K, Hisabori T (2010) CcdA is a thylakoid membrane protein required for the transfer of reducing equivalents from stroma to thylakoid lumen in the higher plant chloroplast. Antioxid. Redox Signal. 13: 1169–1176.

Muthuramalingam M, Seidel T, Laxa M, Nunes de Miranda SM, Gärtner F, Ströher E, Kandlbinder A, Dietz KJ (2009) Multiple redox and non-redox interactions define 2-Cys peroxiredoxin as a regulatory hub in the chloroplast. Mol Plant 2: 1273–88.

Naranjo, B, Mignee C, Krieger-Liszkay A, Hornero-Mendez D, Gallardo-Guerrero L, Cejudo FJ, Lindahl, M (2016) The chloroplast NADPH thioredoxin reductase C, NTRC, controls non-photochemical quenching of light energy and photosynthetic electron transport in Arabidopsis. Plant Cell Environ. 39: 804–822.

Naranjo B, Penzler J-F, Rühle T, Leister D (2021) NTRC Effects on Non-Photochemical Quenching Depends on PGR5. Antioxidants 10:900.

Nikkanen L, Toivola J, Rintamaki E (2016) Crosstalk between chloroplast thioredoxin systems in regulation of photosynthesis. Plant Cell Environ. 39: 1691–1705.

Nikkanen L, Toivola J, Trotta A, Diaz MG, Tikkanen M, Aro EM, Rintamäki E (2018) Regulation of cyclic electron flow by chloroplast NADPH-dependent thioredoxin system. Plant Direct. 2: e00093. doi: 10.1002/pld3.93.

Ojeda V, Perez-Ruiz JM, Gonzalez M, Najera VA, Sahrawy M, Serrato AJ, Geigenberger P, Cejudo FJ (2017a) NADPH thioredoxin reductase C and thioredoxins act concertedly in seedling development. Plant Physiol. 174: 1436–1448.

Ojeda V, Perez-Ruiz JM, Cejudo FJ (2018) 2-Cys peroxiredoxins participate in the oxidation of chloroplast enzymes in the dark. Mol. Plant 11: 1377–1388.

Ojeda V, Pérez-Ruiz JM, González M, Nájera VA, Sahrawy M, Serrato AJ, Geigenberger P, Cejudo FJ (2017b) NADPH Thioredoxin Reductase C and Thioredoxins Act Concertedly in Seedling Development. Plant Physiol 174: 1436–1448.

Okegawa Y, Motohashi K (2015) Chloroplastic thioredoxin m functions as a major regulator of Calvin cycle enzymes during photosynthesis in vivo. Plant J 84: 900–13.

Okegawa Y, Motohashi K (2020) M-Type Thioredoxins Regulate the PGR5/PGRL1-Dependent Pathway by Forming a Disulfide-Linked Complex with PGRL1. Plant Cell 32: 3866–3883.

Okegawa Y, Sato N, Nakakura R, Murai R, Sakamoto W, Motohashi K (2023) x- and y-type thioredoxins maintain redox homeostasis on photosystem I acceptor side under fluctuating light. Plant Physiol :kiad466. doi: 10.1093/plphys/kiad466.

Page ML, Hamel PP, Gabilly ST, Zegzouti H, Perea JV, Alonso JM, Ecker JR, Theg SM, Christensen SK, Merchant S (2004) A homolog of prokaryotic thiol disulfide transporter CcdA is required for the assembly of the cytochrome b6f complex in Arabidopsis chloroplasts. J Biol Chem. 279: 32474–82.

Pan X, Ma J, Su X, Cao P, Chang W, Liu Z, Zhang X, Li M (2018) Structure of the maize photosystem I supercomplex with light-harvesting complexes I and II. Science 360: 1109–1113.

Pesaresi P, Scharfenberg M, Weigel M, Granlund I, Schröder WP, Finazzi G, Rappaport F, Masiero S, Furini A, Jahns P, Leister D (2009) Mutants, overexpressors, and interactors of Arabidopsis plastocyanin isoforms: revised roles of plastocyanin in photosynthetic electron flow and thylakoid redox state. Mol Plant 2: 236–48.

Perez-Ruiz JM, Guinea M, Puerto-Galan L, Cejudo FJ (2014) NADPH thioredoxin reductase C is involved in redox regulation of the Mg-chelatase I subunit in Arabidopsis thaliana chloroplasts. Mol. Plant 7: 1252–1255.

Perez-Ruiz JM, Naranjo B, Ojeda V, Guinea M, Cejudo FJ (2017) NTRC-dependent redox balance of 2-Cys peroxiredoxins is needed for optimal function of the photosynthetic apparatus. Proc. Natl. Acad. Sci. U. S. A. 114: 12069–12074.

Qin X, Suga M, Kuang T, Shen JR (2015) Photosynthesis. Structural basis for energy transfer pathways in the plant PSI-LHCI supercomplex. Science 348: 989–95.

Rey P, Sanz-Barrio R, Innocenti G, Ksas B, Courteille A, Rumeau D, Issakidis-Bourguet E, Farran I (2013) Overexpression of plastidial thioredoxins f and m differentially alters photosynthetic activity and response to oxidative stress in tobacco plants. Front Plant Sci 4: 390.

Richter AS, Peter E, Rothbart M, Schlicke H, Toivola J, Rintamaki E, Grimm B (2013) Posttranslational influence of NADPH-dependent thioredoxin reductase C on enzymes in tetrapyrrole synthesis. Plant Physiol. 162: 63–73.

Serrato AJ, Perez-Ruiz JM, Spinola MC, Cejudo FJ (2004) A novel NADPH thioredoxin reductase, localized in the chloroplast, which deficiency causes hypersensitivity to abiotic stress in Arabidopsis thaliana. J. Biol. Chem. 279: 43821–43827.

Ströher E, Dietz KJ (2008) The dynamic thiol-disulphide redox proteome of the Arabidopsis thaliana chloroplast as revealed by differential electrophoretic mobility. Physiol Plant. 133: 566–83

Takahashi H, Clowez S, Wollman FA, Vallon O, Rappaport F (2013) Cyclic electron flow is redox-controlled but independent of state transition. Nat. Commun. 4: 1954.

Telman W, Liebthal M, Dietz KJ (2020) Redox regulation by peroxiredoxins is linked to their thioredoxin-dependent oxidase function, Photosynth. Res. 145: 31–41.

Thormählen Meitzel T, Groysman J, Ochsner AB, von Roepenack-Lahaye E, Naranjo B, Cejudo FJ, Geigenberger P (2015) Thioredoxin f1 and NADPH-dependen thioredoxin reductase C have overlapping functions in regulating photosynthetic metabolism and plant growth in response to varying light conditions. Plant Physiol 169: 1766–1786.

Thormählen I, Zupok A, Rescher J, Leger J, Weissenberger S, Groysman J, Orwat A, Chatel-Innocenti G, Issakidis-Bourguet E, Armbruster U, Geigenberger P (2017) Thioredoxins play a crucial role in dynamic acclimation of photosynthesis in fluctuating light. Mol. Plant 10: 168–182.

Vaseghi MJ, Chibani K, Telman W, Liebthal MF, Gerken M, Schnitzer H, Mueller SM, Dietz KJ (2018) The chloroplast 2-cysteine peroxiredoxin functions as thioredoxin oxidase in redox regulation of chloroplast metabolism. ELife 7: e38194. DOI: 10.7554/eLife.3819

Wang, P, Liu J, Liu B, Feng D, Da Q, Wang P, Shu S, Su J, Zhang Y, Wang J, Wang HB (2013) Evidence for a role of chloroplastic m-type thioredoxins in the biogenesis of photosystem II in Arabidopsis. Plant Physiol. 163: 1710–1728.

Wolf BC, Isaacson T, Tiwari V, Dangoor I, Mufkadi S, Danon A (2020) Redox regulation of PGRL1 at the onset of low light intensity. Plant J 103: 715–725

